# A SIMPLI (Single-cell Identification from MultiPLexed Images) approach for spatially resolved tissue phenotyping at single-cell resolution

**DOI:** 10.1101/2021.04.01.437886

**Authors:** Michele Bortolomeazzi, Lucia Montorsi, Damjan Temelkovski, Mohamed Reda Keddar, Amelia Acha-Sagredo, Michael J. Pitcher, Gianluca Basso, Luigi Laghi, Manuel Rodriguez-Justo, Jo Spencer, Francesca D. Ciccarelli

**Author notes:** = corresponding author.

## Abstract

Multiplexed imaging technologies enable the study of biological tissues at single-cell resolution while preserving spatial information. Currently, high-dimension imaging data analysis is technology-specific and requires multiple tools, restricting analytical scalability and result reproducibility. Here we present SIMPLI (Single-cell Identification from MultiPlexed Images), a novel, flexible and technology-agnostic software that unifies all steps of multiplexed imaging data analysis. After raw image processing, SIMPLI performs a spatially resolved, single-cell analysis of the tissue slide as wells as cell-independent quantifications of marker expression to investigate features undetectable at the cell level. SIMPLI is highly customisable and can run on desktop computers as well as high-performance computing environments, enabling workflow parallelisation for large datasets. SIMPLI produces multiple tabular and graphical outputs at each step of the analysis. Its containerised implementation and minimum configuration requirements make SIMPLI a portable and reproducible solution for multiplexed imaging data analysis.

SIMPLI is available at: https://github.com/ciccalab/SIMPLI.

## MAIN

A detailed investigation of tissue composition and function in health and disease requires spatially resolved, single-cell approaches that precisely quantify cell types and states as well as their interactions *in situ*. Recent technological advances have enabled to stain histological sections with multiple tagged antibodies that are subsequently detected using fluorescence microscopy or mass spectrometry^1^. High-dimensional imaging approaches such as imaging mass cytometry (IMC)^2^, multiplexed ion beam imaging (MIBI)^3^, co-detection by indexing (CODEX)^4^, multiplexed immunofluorescence (mIF, including cycIF)^5^ and multiplexed immunohistochemistry (mIHC)^6, 7^ enable quantification and localisation of cells in sections from formalin-fixed paraffin-embedded (FFPE) tissues, including clinical diagnostic samples. This is of particular value for mapping the tissue-level characteristics of disease conditions and for predicting the outcome of therapies that depend on the tissue environment, such as cancer immunotherapy. For example, a recent IMC phenotypic screen of breast cancer subtypes revealed the association between the heterogeneity of somatic mutations and that of the tumour microenvironment^8^. Similarly, a CODEX-based profile of FFPE tissue microarrays from high-risk colorectal cancer patients correlated PD1^+^CD4^+^ T cells with patient survival^9^.

The analysis of multiplexed images requires the conversion of pixel intensity data into single-cell data, which can then be characterised phenotypically, quantified comparatively and localised spatially back in the tissue. Currently available tools are technology-specific and cover only some steps of the whole analytical workflow (Table 1). For example, several computational approaches have been developed to process raw images and extract single-cell data either interactively (Ilastik^10^, CellProfiler4^11^, CODEX Toolkit^4^) or via command line (imcyto^12^, ImcSegmentationPipeline^13^). Distinct sets of tools can then perform cell phenotyping (CellProfiler Analyst^14^, Cytomapper^15^, Immunocluster^16^) or analyse cell-cell spatial interactions (CytoMap^17^, ImaCytE^18^, SPIAT^19^, neighbouRhood^20^). Similarly, a few tools enable direct pixel-based analysis through pixel classification^10^ or quantification of pixel positive areas^11^. Despite such a variety of tools, none of them can perform all of the required analytical steps in a common pipeline. Two exception are histoCAT++^21^ and QuPath^22^, which however have been developed specifically for interactive use and are not well suited for the analysis of large datasets. Moreover, all of these tools relies on *ad hoc* configuration files and input formats, making the analysis challenging for users with limited computational skills and restricting the scalability, portability and reproducibility in different computing environments.

**Table 1.**
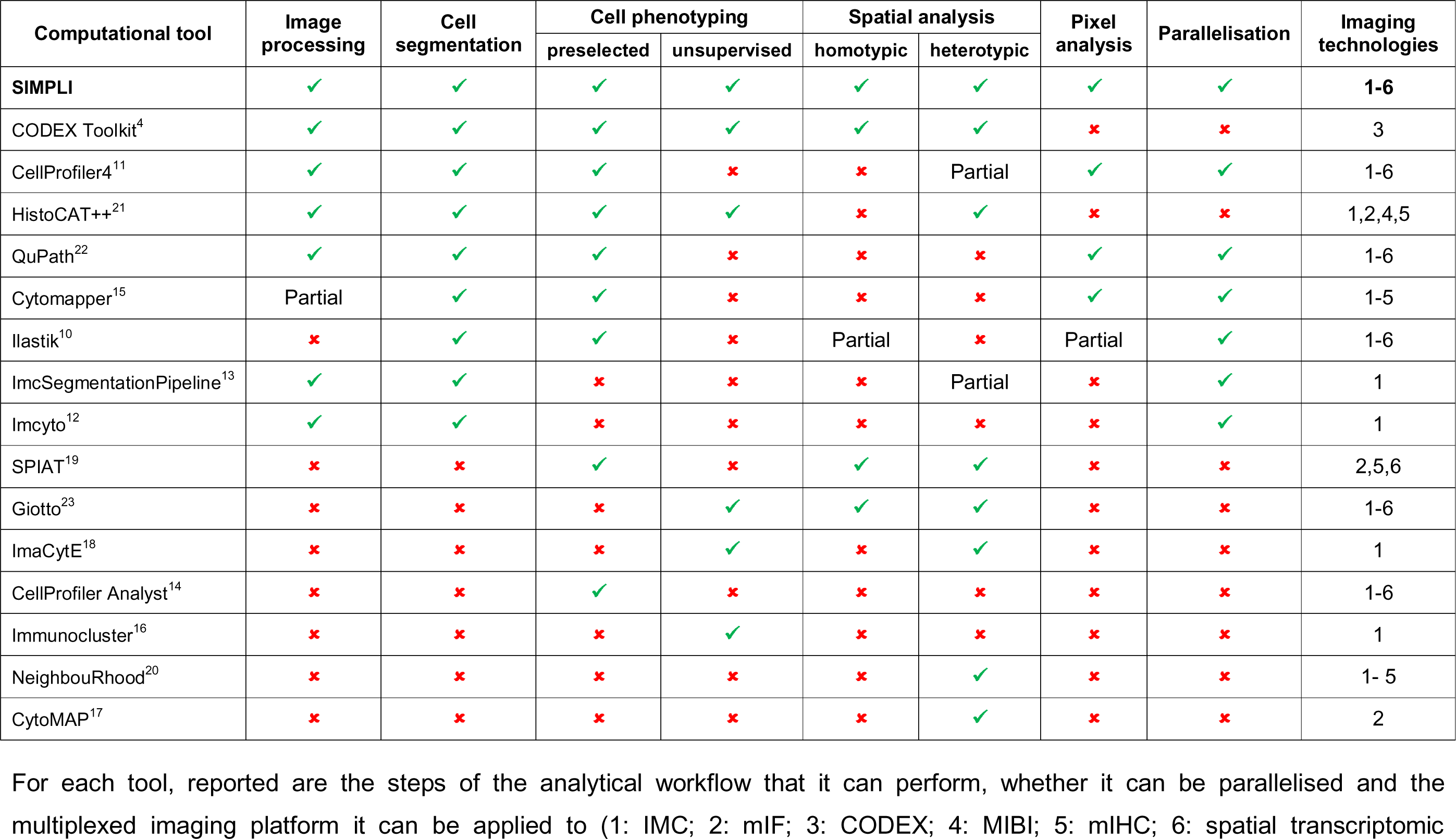
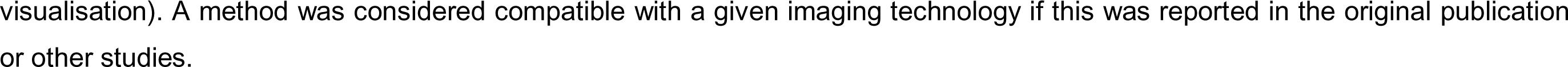
Features of representative tools for the analysis of multiplexed imaging data

Here we introduce SIMPLI (Single-cell Identification from MultiPLexed Images), a tool that combines processing of raw images, extraction of single-cell data, and spatially resolved quantification of cell types or functional states into a single pipeline (Table 1). This is achieved through the integration of well-established tools and newly developed scripts into the same workflow, enabling *ad hoc* configurations of the analysis while ensuring interoperability between its different parts. SIMPLI can be run on desktop computers as well as on high-performance-computing environments, where it can be easily applied to large datasets due to automatic workflow parallelisation. To demonstrate the flexibility of SIMPLI to work with different technologies and experimental conditions, we analyse the phenotypes and spatial distribution of cells in different tissues (human colon, appendix, colorectal cancer) using multiplexed images obtained with distinct technologies (IMC, mIF, CODEX).

## RESULTS

### Overview of the SIMPLI analytical workflow

SIMPLI performs the analysis of multiplexed imaging data in three steps (Methods, Fig. 1), further divided into stand-alone processes (Supplementary Fig. 1). Each process can be run independently or even skipped with the possibility of using alternative input data at each point of the workflow.

**Figure 1.**
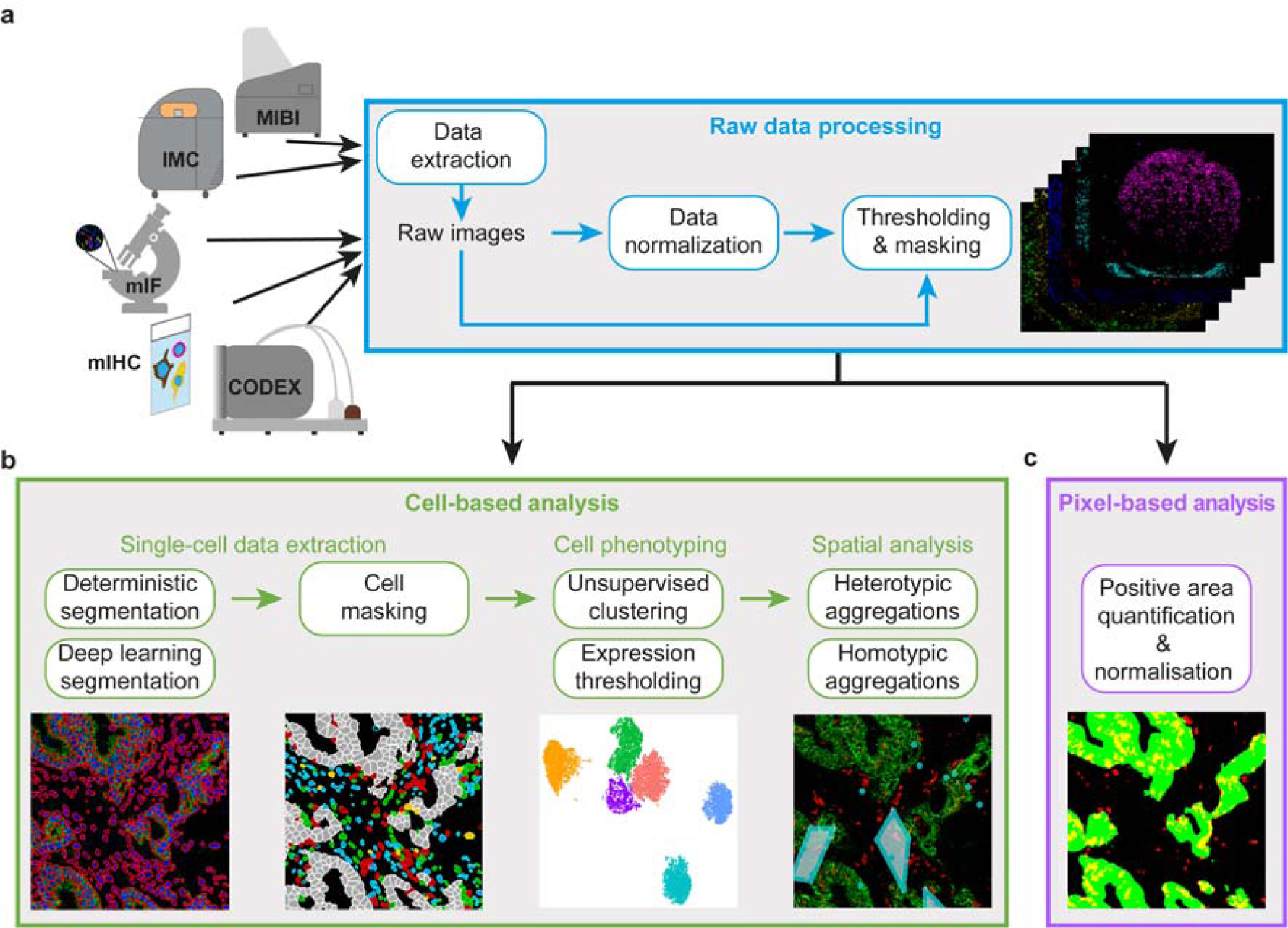
Schematics of the SIMPLI workflow. **a.** Raw images are extracted from IMC or MIBI data or directly imported from other imaging technologies. After their optional normalisation, these images are thresholded to remove the background noise and produce tissue compartment or marker masks. The resulting images can be analysed using a cell-based or a pixel-based approach. **b.** In the cell-based analysis, single cells are segmented with deterministic or deep learning models and phenotyped using unsupervised or supervised approaches. The distribution of cells in the tissue can then be investigated through a spatial analysis of homotypic or heterotypic aggregations. **c.** In the pixel-based approach, areas positive for user-defined combination of markers are measured and normalised over the area of the whole image or of the masks defining compartments or areas positive for certain markers.

The first step of SIMPLI consists of raw data processing (Fig. 1a). Data can be single or multichannel images or text files from a variety of high-dimensional imaging technologies. After data extraction, pixel values for each marker can be normalised by rescaling their values in each sample to allow the user to apply the same thresholds for background noise reduction across samples. Alternatively, sample-specific thresholds can be applied directly to individual, non-normalised images to minimise the effect of non-uniform staining. Finally, masks of specific tissue compartments or markers are derived for subsequent use. The obtained images can then be analysed at the cell (Fig. 1b) and pixel (Fig. 1c) levels.

The cell-based analysis aims to investigate the qualitative and quantitative cell composition of the tissue and is formed of (1) single-cell data extraction, (2) cell phenotyping and (3) spatial analysis of cell-cell distances (Fig. 1b). To extract cell data, SIMPLI implements single-cell segmentation with either a conventional approach based on CellProfiler4^11^ or a deep learning approach based on StarDist^24^. The former enables deterministic filtering based on cells size and shape, as well as marker intensities. The latter applies pre-trained models (either provided by SIMPLI or supplied by the user) to identify cells with high accuracy. After cell segmentation, SIMPLI produces the masks of each individual cell together and calculates the expression values for each marker. Cells belonging to tissue compartments or positive for certain markers can then be identified based on their overlap with previously derived tissue compartment or marker masks. Cell functional phenotypes can be further characterised with two alternative approaches. The first applies unsupervised clustering to all cells or preselected subsets of cells (for example those mapping to specific tissue compartments or positive for certain markers) using marker expression levels. This leads to the unbiased classification of cells into clusters with similar expression profiles indicating similar phenotypes. The second approach identifies cells with designated phenotypes by applying combinations of user-defined thresholds to the expression values of the markers of interest. These thresholds can be identified through an expert guided examination of the original images using the visualisation plots produced by SIMPLI. Finally, a spatial analysis of the distance between cells within the imaged tissue leads to the identification of cell aggregations of the same (homotypic) or different (heterotypic) cell types. In the case of homotypic aggregations, SIMPLI identifies groups of cells of the same type within a user-defined distance and visually localises them as clusters in the tissue image. In the case of heterotypic aggregations, SIMPLI computes the distance distribution between distinct cell types and compares them across cell types and experimental conditions.

The pixel-based approach implemented in SIMPLI enables quantification of areas positive for a specific marker or combination of markers, independently of their association with cells (Fig. 1c). The obtained marker-positive areas are then normalised over the area of the whole image, or those of specific tissue compartments or positive for certain markers using the predefined masks, to allow comparisons across samples. The pixel-based analysis is useful for the investigation of tissue features that are not detectable at the cell level. For instance, extracellular or secreted proteins cannot be quantified with approaches dependent on cell segmentation. In addition, being completely cell agnostic, the pixel-based analysis can provide an independent validation of cell-based observations.

SIMPLI generates tables, plots and images as outputs of each process, thus enabling the visualisation of results at each step of the analysis.

### IMC quantification of secreted and cell-associated IgA in human colon

To test its performance and versatility, we applied SIMPLI to four case studies describing the analysis of multiplexed tissue images of diverse origin, size and resolution and obtained with different technologies (Table 2).

**Table 2.** Description of the case studies used to test SIMPLI.

| Case study | Imaging technology | Analysed samples (n) | Channels (n) | ROI (mm<sup>2</sup>) | Resolution (µm/pixel) | HPC platform | CPU time (h) | Elapsed real time (h) | RAM (GB) | Processes |
| --- | --- | --- | --- | --- | --- | --- | --- | --- | --- | --- |
| 1<br>(Fig.2) | IMC | 6 | 28 | 1.00 | 1.00 | SGE | 00:20:41 | 00:06:10 | 4.1 | • Raw data processing<br>• Cell masking<br>• Single-cell quantification<br>• Pixel intensity comparison |
| 2<br>(Fig.3) | IMC | 1 | 28 | 1.00 | 1.00 | SLURM | 00:06:25 | 00:05:30 | 4.2 | • Raw data processing<br>• Cell masking<br>• Unsupervised clustering<br>• Expression thresholding<br>• Homotypic cell distances |
| 3<br>(Fig.4) | mIF | 1 | 7 | 5.45 | 0.50 | SLURM | 00:11:45 | 00:08:23 | 16.7 | • Thresholding & masking<br>• Expression thresholding<br>• Heterotypic cell distances |
| 4<br>(Fig.5) | CODEX | 35 | 58 | 1.13 | 0.38 | SGE | 02:32:35 | 00:26:01 | 22.5 | • Expression thresholding<br>• Heterotypic cell distances |

For each case study, listed are the imaging technologies used to generate the tissue images, the number of samples and markers used, the size of the analysed region of interest (ROI), the resolution of the obtained images, the high performance (HPC) platform and the computational resources employed to perform the analysis. These include the central processing unit (CPU) time and the elapsed real time, as well as the maximum random access memory (RAM) memory used. Finally, the specific analytical processes performed in each case study are also listed (single cell segmentation was performed in all of them). SGE: Sun Grid Engine; SLURM: Simple Linux Utility for Resource Management.

As a first case study, we used SIMPLI to compare the levels of secreted and cell-associated immunoglobulin A (IgA), the major immunoglobulin isotype in intestinal mucosa^25^, from IMC-derived multiplexed images of normal human colon. We stained six colon sections (CLN1-CLN6, Supplementary Table 1) with 26 antibodies marking T cells, macrophages, dendritic cells and B cells as well as stromal components (Supplementary Table 2) and ablated one region of interest (ROI) per sample.

Using SIMPLI, we extracted and normalised the 28 single channel images (26 antibodies and two DNA intercalators) for each of the six ROIs and combined them into a single image per ROI (Fig. 2a). This normalisation enabled selection of a single threshold for each marker to be used across all samples, thus reducing the complexity of the analysis configuration. By applying these thresholds to the E-cadherin and vimentin expression, we obtained the masks for the epithelium and the lamina propria, respectively (Fig. 2b). We used these masks to assign cells to the two compartments and normalise marker values or positive areas in the downstream analyses.

**Figure 2.**
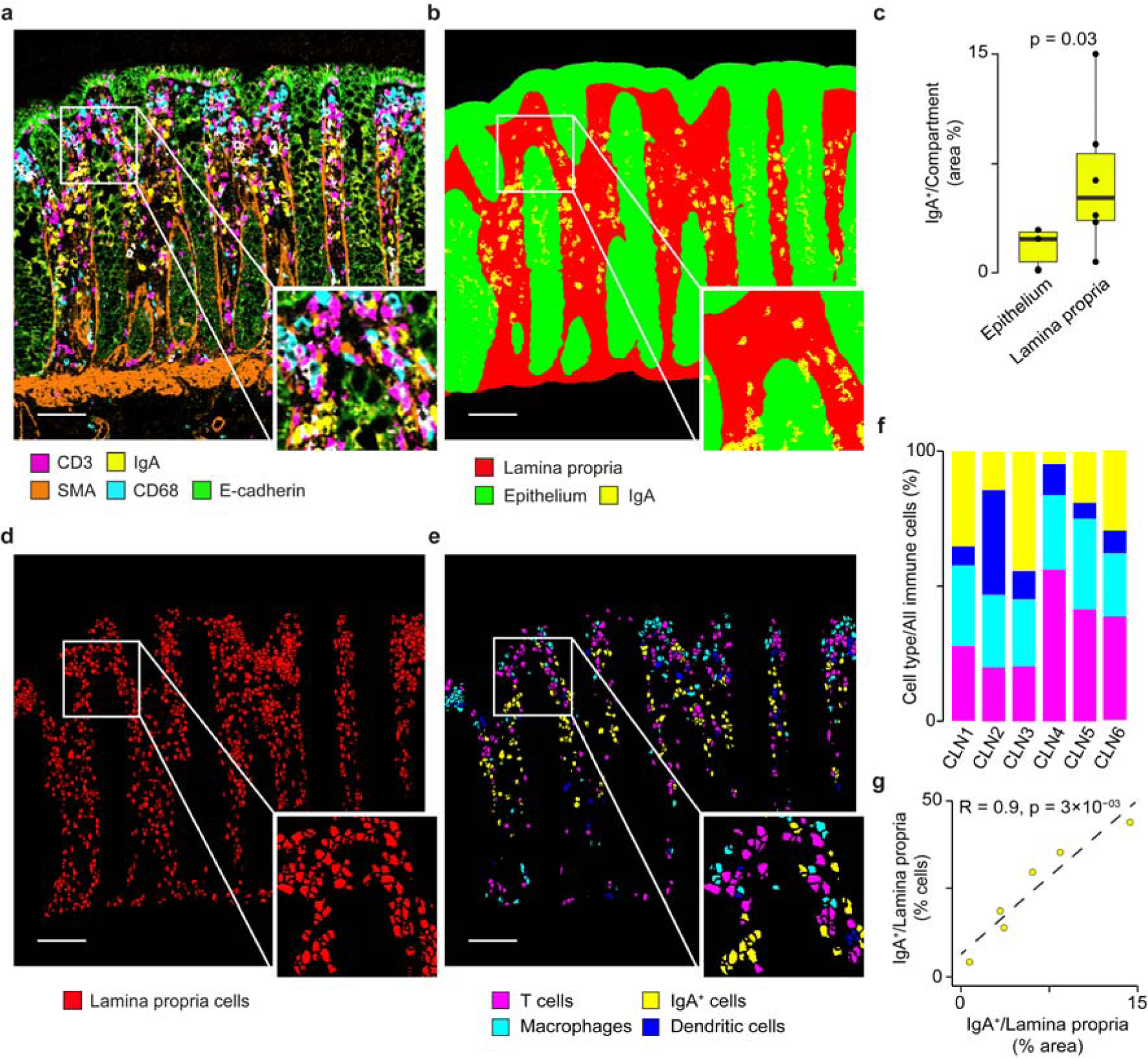
IgA quantification in human colon mucosa. **a.** IMC image of a representative sample (CLN6) of normal colon mucosa after extraction and normalisation of raw data. **b.** Masks defining the lamina propria and the epithelial compartments overlaid with IgA^+^ areas. Lamina propria and epithelial masks were obtained by thresholding the vimentin and E-cadherin channels, respectively. **c.** Comparison of normalised IgA^+^ areas in the lamina propria and epithelial compartments in CLN1-CLN6. Normalised areas were measured as the proportion of IgA^+^ area over the lamina propria and epithelium masks, respectively. Distributions were compared using a two-sided Wilcoxon test. **d.** Outlines of the cells in the lamina propria. After single-cell segmentation, all cells overlapping with the lamina propria mask by at least 30% of their area were considered as cells resident in the lamina propria. **e.** Outlines of immune cells resident in the lamina propria identified according to the highest overlap between their area and the masks for IgA^+^ cells, T cells, macrophages and dendritic cells. **f.** Relative proportions of T cells, IgA^+^ cells, macrophages and dendritic cells over all immune cells in the lamina propria across CLN1-CLN6. **g.** Correlation between normalised IgA^+^ area and the proportion of IgA+ cells over the total immune cells in in the lamina propria in CLN1-CLN6. Pearson correlation coefficient R and associated p-value are shown. Scale bar in all images = 100μm.

We then used the pixel-based approach to quantify both the IgA expressed by the plasma cells resident in the diffuse lymphoid tissue of the lamina propria as well as the secreted IgA undergoing transcytosis to traverse the epithelial compartment (Fig. 2b). As expected, most secreted IgA was localised in the epithelial crypts with only minimal presence of IgA^+^ area in the surface epithelium (Supplementary Fig. 2a). Quantification of the normalised IgA^+^ areas in the two compartments (Supplementary Fig. 2b) confirmed higher IgA^+^ levels in the lamina propria than in the epithelium (Fig. 2c). To assess the impact of image normalisation, we repeated the same analysis starting from the raw images and applying sample-specific thresholds to remove the background noise. The resulting IgA levels correlated linearly with those obtained from normalised images (Supplementary Fig. 2c), showing that data normalisation has no impact on the results,

Next, we quantified the IgA^+^ plasma cells in the lamina propria using the cell-based approach. First, we performed single-cell segmentation with the deterministic approach and retained only cells overlapping for at least 30% or their area with the lamina propria mask (Fig. 2d). We verified that varying the threshold of the overall had a minimal impact on the proportion of cells assigned to the lamina propria (Supplementary Fig. 2e). We then identified IgA^+^ plasma cells, T cells, macrophages, and dendritic cells resident in the lamina propria according to the highest overlap between the cell area and the mask of each immune cell population (Fig. 2e). Again, we verified that the relative proportion of these cell populations changed only minimally varying the threshold of the overlap with the lamina propria mask (Supplementary Fig. 2e). Finally, we quantified the four immune cell populations across the six samples and observed that IgA^+^ plasma cells constitute approximately 25% of all identified immune cells (Fig. 2f). This is consistent with previous quantifications of the fraction of plasma cells over the total mononucleated cells in the lamina propria of healthy individuals^26^.

The relative proportion of IgA^+^ plasma cells positively correlated with the normalised IgA^+^ area in the lamina propria, demonstrating that the quantification from the single-cell analysis is supported by the cell agnostic measurements at the pixel-level (Fig. 2g).

### Localisation of T follicular helper cells in IMC images of a germinal centre

As a second case study, we used SIMPLI to localise the immune cell populations within a FFPE section of healthy human appendix (APP1, Supplementary Table 1). After staining the tissue section with the 28 markers (26 antibodies and two DNA intercalators) used previously (Supplementary Table 2), we performed IMC and used SIMPLI to extract and normalise the single channel images from the raw IMC data for the ROI. The resulting combined image revealed a germinal centre in the B cell area and follicle-associated epithelium forming the boundary with the appendiceal lumen (Fig.3a).

We performed single-cell segmentation with both approaches implemented in SIMPLI and observed high overlap (Supplementary Fig. 3a), indicating good concordance between the two methods. We then classified the 7573 cells from CellProfiler4 in immune and epithelial cells based on the highest overlap with the corresponding masks obtained in the data extraction step (Fig. 3b). Using both phenotyping approaches available in SIMPLI, we characterised the phenotypes of T cells, that overall constituted approximately 27% of all cells (Fig. 3c). First, we applied unsupervised clustering using seven markers of T cell function (Supplementary Table 2). After inspection of the resulting clusters at different resolution levels, we selected 0.25 resolution which returned five distinct cell clusters (Fig. 3d). Based on the marker expression profiles, we assigned cluster 1 to CD4^+^ T cells, cluster 2 to CD8^+^CD45RO^+^ T cells, cluster 3 to CD4^+^CD45RA^+^ T cells, cluster 4 to CD4^+^CD45RO^+^ T cells and cluster 5 to CD4^+^PD1^+^ T cells (Fig. 3e). The latter likely represented a set of PD1^+^ T follicular helper cells known to be located in the germinal centre^27^. Interestingly, at higher resolution, cluster 5 was further divided into two smaller clusters showing PD1 high and low expression (Supplementary Figure 3a). Similarly, clusters 1 and 2 were further divided into smaller subpopulation based on CD4 and CD45RO expression levels, respectively (Supplementary Figure 3a).

**Figure 3.**
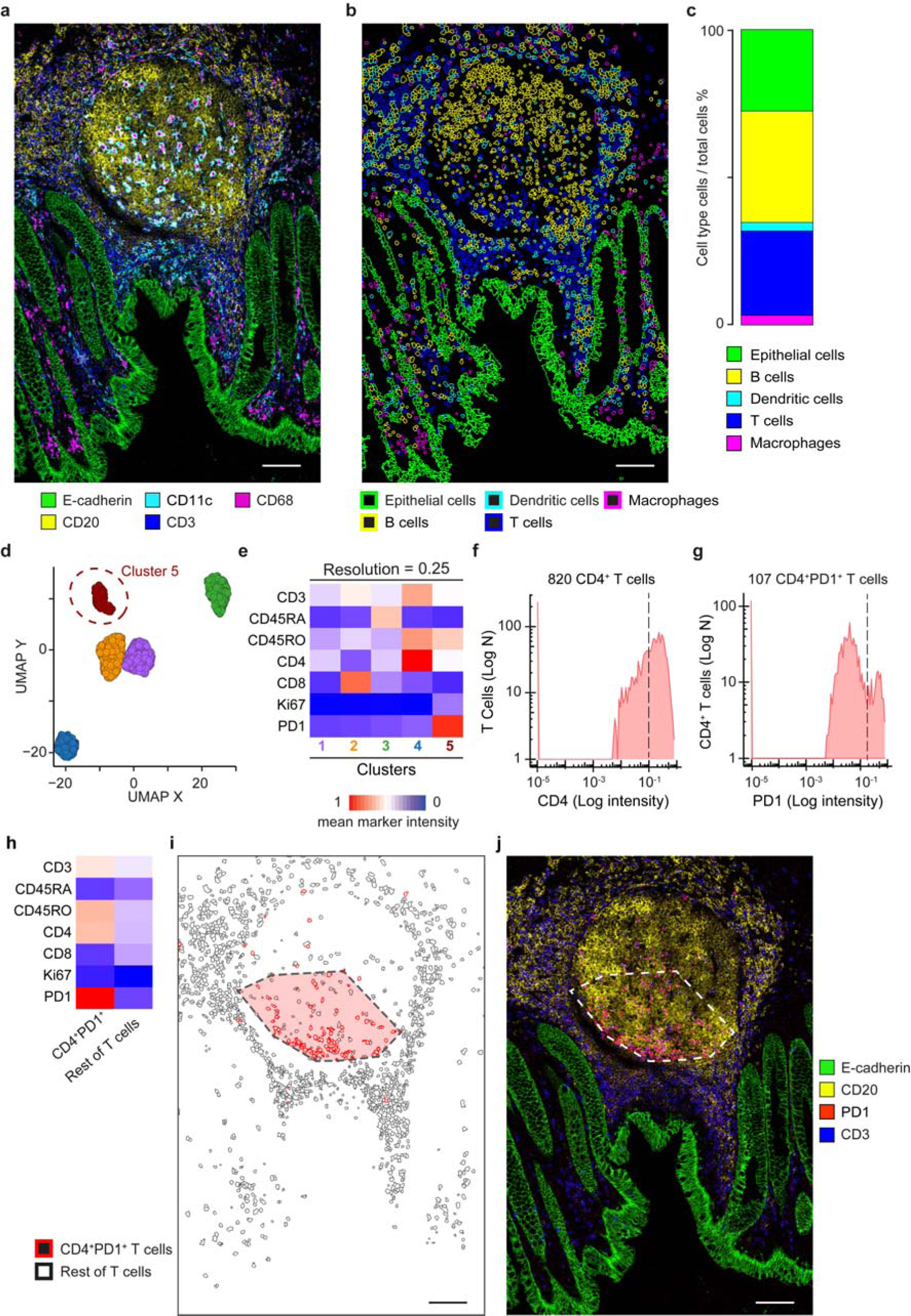
Single-cell characterisation of T cells in a human germinal centre. **a.** IMC image of a normal appendix (APP1) showing a central germinal centre with the columnar epithelium delimiting the appendiceal lumen. **b.** Outlines of T cells, B cells, macrophages, dendritic and epithelial cells identified through the highest overlap with the respective masks. **c.** Proportions of T cells, B cells, macrophages, dendritic and epithelial cells over all cells. **d.** UMAP plot of 1,466 T cells grouped in five clusters resulting from unsupervised clustering according to the expression of seven markers of T cell function (Supplementary Table 2). Cluster 5 (circled) corresponds to PD1^+^CD4^+^ T cells. **e.** Expression profiles of the five clusters identified in (d). The mean intensity value of each marker across all cells is reported. The colour scale was normalized across all markers and cells. **f.** Density plots of CD4 expression in T Cells. Cells with ≥0.1 CD4 expression were considered as CD4^+^ T cells. **g.** Density plot of PD1 expression in CD4^+^ T cells. Cells with ≥0.15 PD1^+^ expression were considered as PD1^+^CD4^+^ T cells. Thresholds for CD4 and PDL1 were identified through histological inspection of the PD1 channel images. **h.** Expression profiles of the PD1^+^CD4^+^ T cells and rest of T cells. For both populations, the mean intensity value of each marker across all cells is shown. The colour scale was normalized across all markers and cells. **i.** Position map of T cells within the ROI. The area of a high-density cluster of ≥5 PD1^+^CD4^+^ T cells per 10,000 *μ*m^2^ is highlighted in red. **j.** IMC image showing the localisation of the PD1 signal within the ROI. Scale bar for all images = 100μm.

We re-identified these PD1^+^ T follicular helper cells with the second phenotyping approach based on expression thresholding of CD4 and PD1 (Fig.1b). Starting from all T cells, we first extracted CD4^+^ T cells (≥0.1 CD4 expression, Fig. 3f) and, within those, we further identified PD1^+^ cells (≥0.15 PD1 expression, Fig. 3g). Both thresholds were chosen after manual inspection of the histological images. The expression profile of the resulting PD1^+^CD4^+^ T cells (Fig 3h) closely recapitulated that of cluster 5 (Fig. 3e). We repeated the same analysis for clusters 1 to 4 confirming the high overlap between cells in unsupervised clusters and those re-identified using marker expression thresholds (Supplementary Fig. 3b). Moreover, these cells showed similar expression profiles (Supplementary Fig. 3c) and spatial localisation (Supplementary Fig. 3d), indicating that cell phenotypes identified with unsupervised clustering can be confirmed through user-guided thresholding of marker expression.

Finally, we investigated the spatial localisation of PD1^+^ T follicular helper cells within the ROI by analysing their homotypic aggregations. This allowed us to localise a single high-density cluster containing 84% of PD1^+^CD4^+^ T cells within the germinal centre (Fig. 3i). This distribution of PD1^+^CD4^+^ T cells was in accordance with the localisation of T helper cells in the in the follicles of secondary lymphoid organs^27^ and was confirmed by the histological inspection of the tissue image (Fig. 3j).

### mIF analysis of spatially resolved cell-cell interactions in rectal cancer

As a third case study, we applied SIMPLI to the spatial analysis of mIF-derived images of a rectal cancer sample (CRC1, Supplementary Table 1) stained with anti CD8, PD1, Ki67, PDL1, CD68, GzB and DAPI antibodies (Supplementary Table 2). We focused on a 5mm^2^ ROI that was rich in T cells at the invasive margins of the tumour (Fig. 4a). This allowed us to characterise the cell-cell interactions between PDL1^+^ cells and PD1^+^CD8^+^ T cells at the tumour boundary in a larger ROI, supporting the scalability of SIMPLI to the analysis of large regions (Table 2).

**Figure 4.**
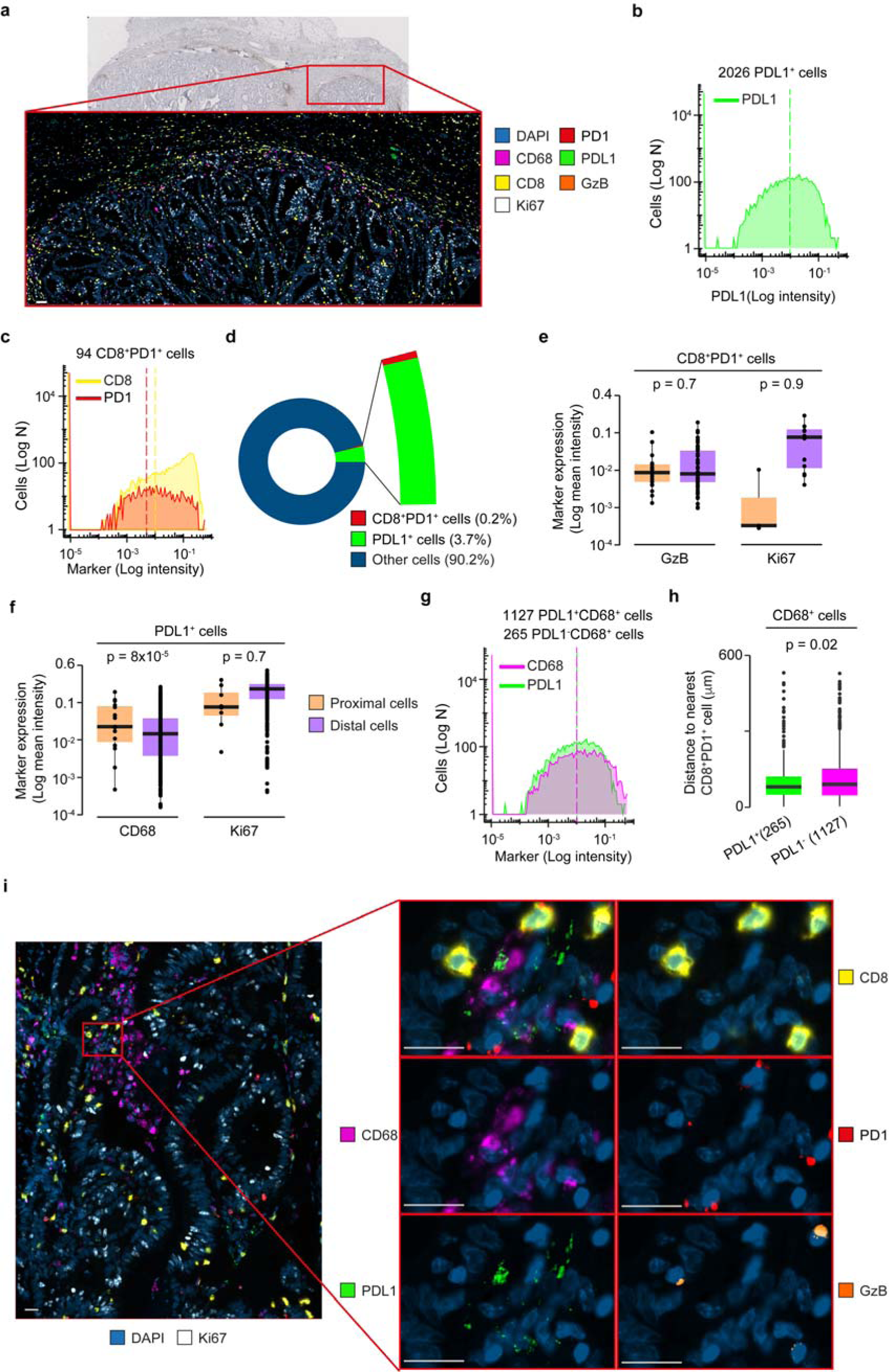
Characterisation of PDL1^+^ and PD1^+^ cells at the tumour invasive margins. **a.** CD3 immunohistochemistry (main image) and sequential mIF image (zoom-in, 20x magnification) of a rectal cancer sample (CRC1). The mIF image corresponded to a 5mm^2^ tissue area at the invasive margins of the tumour and was obtained by combining the pre-processed images of seven markers. Scale bar = 50µm. **b.** Density plot of PDL1 expression in CD8^-^ cells. Cells with ≥0.01 PDL1 expression were considered as PDL1^+^ cells. **c.** Density plots of CD8 and PD1 expression in T cells. Cells with ≥0.01 expression and ≥ 0.005 PD1 expression were considered as PD1^+^ CD8^+^ T cells. Expression thresholds were identified through histological inspection of PDL1, CD8 and PD1 channel images and are indicated as dotted lines in the corresponding plots. **d.** Proportions of PD1^+^CD8^+^ cells, PD1^-^CD8^+^ T cells and PDL1^+^ cells over total cells. **e.** Comparison of the mean intensity of GzB and Ki67 between PD1^+^CD8^+^ T cells proximal and distal to PDL1^+^cells. Proximal PD1^+^CD8^+^ T cells were defined as those at less than 12μm from a PDL1^+^ cell. **f.** Comparison of the mean intensity of CD68 and Ki67 between PDL1^+^cells proximal and distal to PD1^+^CD8^+^ T cells. Proximal PDL1^+^ cells were defined as those at less than 12μm from a PD1^+^CD8^+^ T cell. Distributions in (**e**) and (**f**) were compared using a two-sided Wilcoxon test. **g.** Density plots of CD68 and PD1 expression in all cells. Cells with ≥0.01 CD68 and PDL1 expression were considered as PDL1^+^CD68^+^ cells. **h.** Comparison of distance of PDL1^+^ and PDL1^-^CD68^+^ macrophages to the nearest PD1^+^CD8^+^ T cell. **i.** High resolution (40x magnification) mIF image of PD1^+^CD8^+^ T cells in close proximity to PDL1^+^CD68^+^ cells. Zoom in images show each marker separately and merged. Scale bar = 20µm. All distributions were compared using a two-sided Wilcoxon test.

After image pre-processing and single-cell segmentation, we identified PDL1^+^ and PD1^+^CD8^+^ cells by applying expert-defined thresholds to PDL1 (≥0.01), CD8 (≥0.01), and PD1 (≥0.005) expression levels, respectively. We extracted 2026 PDL1^+^ cells (Fig. 4b) and 3177 CD8^+^ cells, 94 of which also expressed PD1 (Fig. 4c). The two sets of PDL1^+^ and PD1^+^CD8^+^ cells constituted 3.7% and 0.2% of all cells in the analysed region, respectively (Fig. 4d).

We characterised the spatial relationship between these cells, focusing on the ones in close proximity to each other. Using the Euclidean distances between their centroids, we identified 35 PDL1^+^ cells and 21 PD1^+^CD8^+^ T cells at a distance lower than 12μm apart, which corresponded to twice the maximum cell radius length. We considered these cells proximal enough to be engaging in PD1-PDL1 mediated interactions. By comparing PD1^+^CD8^+^ T cells proximal to PDL1^+^ cells and PD1^+^CD8^+^ T cells distal to PDL1^+^ cells, we found no difference in the expression of cytotoxicity (GzB) or proliferation (ki67) markers (Fig. 4e). This is in line with the broad range of cytotoxic activity in this T cell subset observed in colorectal cancer^28^. On the contrary, PDL1^+^ cells proximal to PD1^+^CD8^+^ T cells expressed higher levels of CD68 than PDL1^+^ cells distal to PD1^+^CD8^+^ T cells (Fig. 4f), suggesting spatial proximity between PDL1^+^ macrophages and PD1^+^CD8^+^ T cells. To validate this observation, we identified 1392 macrophages by applying an expert-defined threshold to CD68 expression (≥0.01, Fig. 4g) and classified them as PDL1^-^ and PDL1^+^ cells, respectively using 0.1 PDL1 expression threshold. Comparing the distance of the resulting two populations from the nearest PD1^+^CD8^+^ T cells, we confirmed that PDL1^+^CD68^+^ macrophages were significantly closer to PD1^+^CD8^+^ T cells than PDL1^-^ CD68^+^ macrophages (Fig. 4h). By inspecting the imaged tissue at 40x magnification, we confirmed the localisation of PDL1^+^CD68^+^ macrophages in close proximity to PD1^+^CD8^+^ cells, as well as the presence of both PD1^+^CD8^+^GzB^-^ T cells and PD1^+^CD8^+^GzB^+^ T cells proximal to PDL1^+^ cells (Fig. 4i).

### Comparison of cell distances in CODEX images of colorectal cancer subtypes

As a fourth case study, we used SIMPLI to compare the distances between immune cells and tumour or endothelial cells in CLR (Crohn’s-like reaction) and DII (diffuse inflammatory infiltration) colorectal cancer subtypes^9^. The high-dimensional imaging data derived from 35 colorectal cancer samples (Supplementary Table 1) and were obtained using CODEX with a 56 marker panel^9^ (Supplementary Table 2). Such a large number of antibodies enabled the identification and spatial localisation of T cells, B cells, plasma cells, macrophages, NK cells, granulocytes, dendritic cells, tumour cells, neuroendocrine cells, smooth muscle, nerves, lymphatic and blood vessels (Fig 5a).

**Figure 5.**
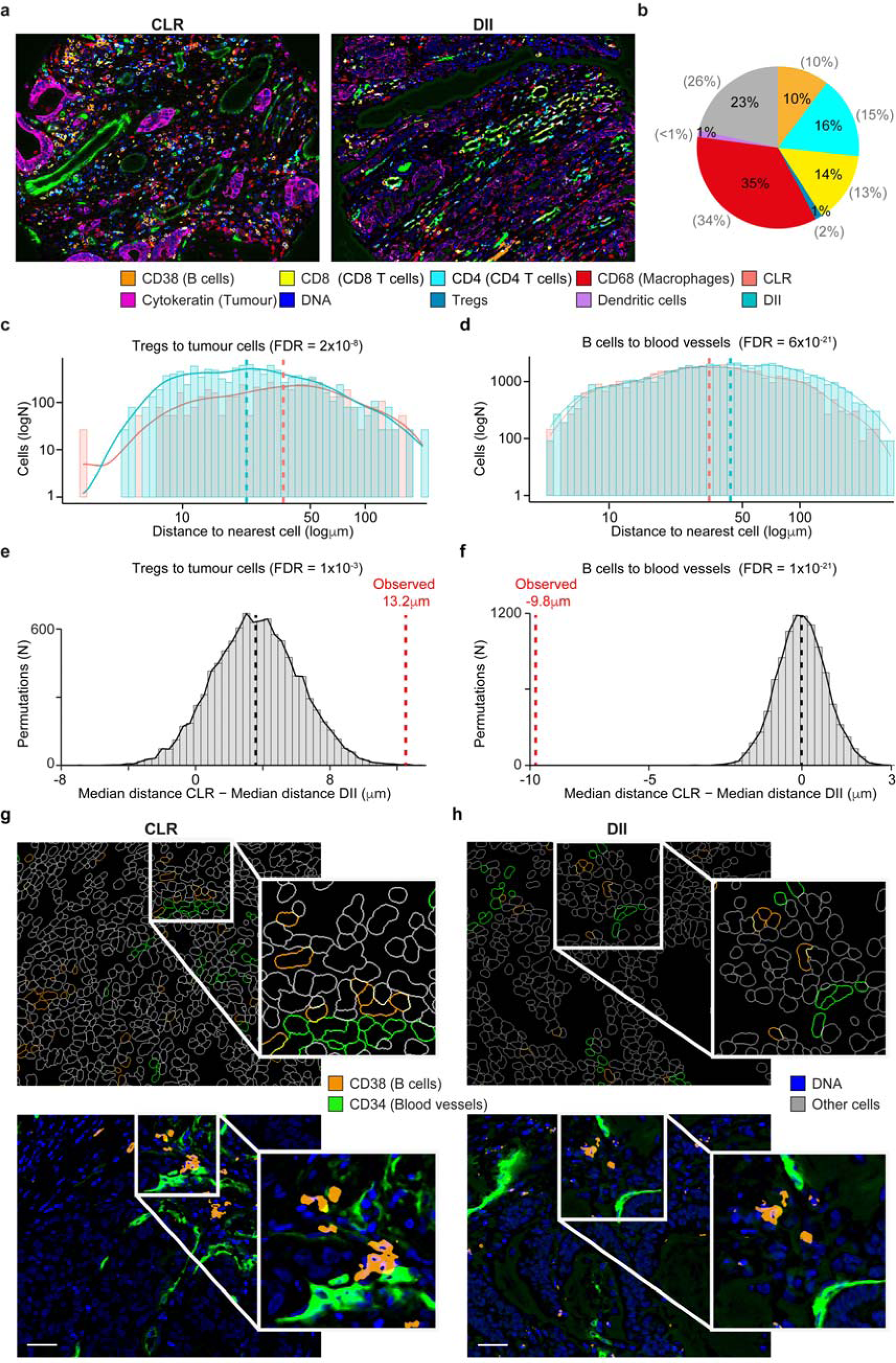
Spatial localisation of immune cells in two colorectal cancer subtypes. **a.** CODEX images of two representative CLR (CRC_12_24) and DII (CRC_31_16) colorectal cancer samples. **b.** Proportions of CD8^+^ T cells, CD4^+^ T cells, Tregs, macrophages, dendritic cells, B cells and other mixed immune cell populations across the 35 analysed samples. Cell types were identified by applying expert-defined thresholds to the expression intensity of representative markers and normalised over the total non-cancer cells. These thresholds were derived through histological inspection of the channel images. The cell proportion corresponding to each population from the original study^9^ is reported in brackets. Distance distribution of Tregs to the nearest tumour cell (**c**) and of B cells to the nearest endothelial cell (**d**) of CLR and DII samples. Distances between cell pairs were calculated using the cell centroids coordinates and the resulting distributions were compared between CRC subtypes using a two-sided Wilcoxon test. Benjamini-Hochberg FDR correction was applied for testing over ten cell type comparisons. Only differences of at least 8µm and with FDR <0.1 were considered significant. Dashed lines represent the medians of the distributions.Distribution of the expected differences between the median distances of Tregs to the nearest tumour cell (**e**) and of B cells to the nearest endothelial cell (**f**) in CLR and DII samples. Expected values were calculated with a permutation test, where cell identities were randomly reassigned for 10000 times within each sample. The resulting median values were compared to the observed differences with a two-tailed permutation test adjusted for multiple hypothesis testing with the Benjamini-Hochberg correction. Single-cell outlines of B cells and blood vessels (upper panel) and associated images (lower panel) form a representative CLR (CRC_17_34) **(g)** and DII (CRC_15_29) **(h)** sample. Scale bar = 100μm.

After single-cell segmentation, we quantified the main cell types identified in the original study^9^ by applying expert-defined thresholds to the expression of markers representative of each population (CDX2, MUC1 or cytokeratin for tumour cells; CD34 or CD31 for endothelial cells; vimentin for stromal cells; CD11c for dendritic cells; CD38 for B cells; CD3 and CD4 for CD4^+^ T cells; CD3, CD4 and FOXP3 for Tregs; CD3 and CD8 for CD8^+^ T cells, CD68 for macrophages). The obtained relative proportions of immune cells across all samples were highly concordant with those reported the original study^9^ (Fig 5b).

We then measured the distances of the main immune cell types from tumour cells and blood vessels by performing a heterotypic spatial analysis. First, we calculated the distances of each macrophage, CD8^+^ T cell, CD4^+^ T cell, Treg and B cell to the nearest tumour cell or endothelial cell using the coordinates of the cell centroids. From these, we derived the corresponding distance distributions from the nearest tumour cell or endothelial cell in each sample. Finally, we compared the resulting distributions between CLR and DII colorectal cancer subtypes. After correcting for multiple testing, we considered biologically relevant only differences between the median distances of the two subtypes bigger than 8µm, corresponding to the diameter of B and T lymphocytes^29^. With this approach, we found that Tregs were significantly closer to tumour cells in DII (median distance = 22.4µm) compared to CLR (35.6µm, Fig 5c). On the contrary, B cells were more proximal to blood vessels in CLR (33.5µm) than in DII (43.3µm, Fig 5d). We further supported these results with a permutation test, where we re-labelled randomly the identities to all cells in each sample for 10000 times to derive an expected distribution of differences in distances between CLR and DII cells. The comparisons of observed values to the expected distributions, confirmed that Tregs were significantly closer to tumour cells in DII (Fig.5e) while B cells were more proximal to blood vessels in CLR (Fig 5f). We supported the distinct spatial distributions of B cells in CLRs (Fig 5g) and DII (Fig 5h) through independent histological image inspection.

This novel result, not reported in the original study, showcases the discovery potential of the quantitative analysis of spatial relationships between cell populations implemented in SIMPLI. Additionally, the SIMPLI graphical representations of the tissue composition as an overlay of cell boundaries color-coded by cell populations greatly facilitates the visual inspection of their spatial interactions in their original tissue context.

## DISCUSSION

SIMPLI is an open-source, customisable and technology-independent tool for the analysis of multiplexed imaging data. It enables the processing of raw images, the extraction of cell data and the spatially resolved quantification of cell types or functional states as well as a cell-independent analysis of tissues at the pixel level, all within a single platform (Table 1).

In comparison to currently available software, SIMPLI increases the portability, scalability and reproducibility of the analysis (Table 2). Moreover, it can easily accommodate specific analytical requirements across a wide range of tissues and imaging technologies at different levels of resolution and multiplexing through user-friendly configuration files. SIMPLI interoperates with multiple software and programming languages by leveraging workflow management and containerisation. This makes the inclusion of new algorithms, features and imaging data formats easy to implement, such as alternative methods of cell-segmentation, pixel and cell classification or a Graphical User Interface for interactive data visualisation

Multiplexed imaging methods have proven to be a powerful approach for the study of tissues through the in-depth characterisation of cell phenotypes and interactions. SIMPLI represents an effort to make these analyses more accessible to a wider community. This will enable exploitation of highly multiplexed imaging technologies for multiple applications, ranging from basic life science and pharmaceutical research to precision medical use in the clinics.

## ONLINE METHODS

### SIMPLI description and implementation

SIMPLI’s workflow is divided into three steps (raw image processing; cell-based analysis; pixel-based analysis), which are constituted of multiple stand-alone processes (Fig. 1 and Supplementary Fig. S1). Processes can be executed sequentially or independently from the command-line or through a configuration file that can be edited with any text editor. This allows the user to skip some of them and use alternative input data for downstream analyses. Additionally, parameters and options can be specified through the same configuration files without the need to setup tool-specific input files in any specific directory structure.

Raw data from IMC or MIBI experiments (.mcd or .txt files) are converted into single or multi-channel .tiff images with imctools^30^. Data from other multiplexed imaging platforms may be supplied directly as raw single or multi-channel tiff images (Supplementary Fig. 1a). Raw images can be thresholded individually to minimise the effect of non-uniform staining and then used directly for the cell- and pixel-based analyses. Alternatively, they can be first normalised across samples by rescaling pixel values of each channel up to the 99^th^ percentile of the distribution using the EBImage^31^ package and custom R scripts. Normalised images can then be processed with CellProfiler4^11^ to generate thresholded images and masks of tissue compartments or markers to be used in the following steps.

Pixel-based and cell-based analyses can be run as single workflows or in parallel within the same run. Both of them provide multiple outputs of the various processes, including tabular text files, visualisation plots and comparisons across samples (Supplementary Fig. 1).

The cell-based analysis is composed of cell data extraction, cell phenotyping and spatial analysis (Supplementary Fig. 1b). The extraction of cell data starts with single-cell segmentation using CellProfiler4^11^ or StarDist^24^ with scikit-image^32^ used for feature extraction. In the latter case, default models or user-provided trained models can be used. Cell segmentation returns (1) single-cell data consisting of the marker expression values and the coordinates of each cell in the ROI and (2) the ROI segmentation mask marking all the pixels belonging to each cell with its unique identifier. Cells mapping to tissue compartments or positive for certain markers can then be identified based on their overlap with the tissue compartments or marker masks derived in the previous step. These cells are visualised in the ROI as outlines, while their proportions are quantified in barplots and boxplots.

All cells, or only those in specific tissue compartments or positive for certain markers can be further phenotyped using two approaches. The first consists of unsupervised clustering based on the marker expression values using Seurat^33^. Cells are represented as nodes in a k-nearest neighbour graph based on their Euclidean distances in a principal component analysis space. This graph is then partitioned into clusters using the Louvain algorithm^34^ at user-defined levels of resolution leading to the unsupervised identification of cell phenotypes. Clusters of cell phenotypes are plotted as scatterplots in Uniform Manifold Approximation and Projection (UMAP)^35^ space. The second phenotyping approach is based on user-defined thresholds of marker expression values that can be combined using logical operators for the identification of designated cell phenotypes. The distributions of cells are represented as density plots based on the marker expression levels. In both phenotyping approaches, the expression profiles of the cell types are plotted as heatmaps, their proportions quantified in barplots and boxplots and their locations in the ROI visualised as cell outlines.

Once cell populations and phenotypes have been identified, the spatial analysis investigates the distance between cells of the same (homotypic aggregations) or different (heterotypic aggregations) types. In the homotypic analysis, clusters of cells of the same type within a user-defined distance are identified with DBSCAN^36^ as implemented in the fpc^37^ R package. These homotypic cell aggregations are visualised as position maps, reporting cell location and high-density clusters in the ROI. In the heterotypic analysis, the cell distances, defined as the Euclidean distances between cell centroids, are computed using custom R scripts and visualised as density plots. The homotypic and heterotypic spatial analyses can be run in parallel or singularly on one or more sets of cells.

The pixel-based analysis quantifies areas positive for user-defined combination of markers using the EBImage^31^ package with custom R scripts (Supplementary Fig. 1c). These measurements are performed starting from the thresholded images produced in the raw image processing step (Supplementary Fig. 1a). The marker positive areas obtained in this way are then normalised over the area of the whole image or specific tissue or marker compartments. The resulting normalised positive areas can then be quantified in barplots and boxplots.

SIMPLI is implemented as a Nextflow^38^ pipeline employing Singularity containers^39^ hosted on Singularity Hub^40^ to manage all the libraries and software tools. This allows SIMPLI to automatically manage all dependencies, irrespective of the running platform. Nextflow also manages automatic parallelisation of all processes while still allowing the selection of parts of the analysis to execute.

### Sample description

Six FFPE blocks of normal (non-cancerous) colon mucosa (CLN1-CLN6), one of normal appendix (APP1), and one of rectal cancer (CRC1) were obtained from eight individuals who underwent surgery for the removal of colorectal cancers (Supplementary Table 1). All blocks were reviewed by an expert pathologist (MRJ). All patients provided written informed consent in accordance with approved institutional guidelines (University College London Hospital, REC Reference: 20/YH/0088; Istituto Clinico Humanitas, REC Reference: ICH-25-09).

### Staining and IMC ablation of human colon mucosa and appendix

Four µm-thick sections were cut from each block of samples CLN1-CLN6 and APP1 with a microtome and used for staining with a panel of 26 antibodies targeting the main immune, stromal and epithelial cell populations of the gastrointestinal tract (Supplementary Table 2). The optimal dilution of each antibody in the panel was identified by staining and ablating FFPE appendix sections. The resulting images were reviewed by a mucosal immunologist (J.S.) and the dilution giving the best signal to background ratio was selected for each antibody (Supplementary Table 2). To perform the staining for IMC, slides were dewaxed after a one-hour incubation at 60°C, rehydrated and heat-induced antigen retrieval was performed with a pressure cooker in Antigen Retrieval Reagent-Basic (R&D Systems). Slides were incubated in a 10% BSA (Sigma), 0.1% Tween (Sigma), and 2% Kiovig (Shire Pharmaceuticals) Superblock Blocking Buffer (Thermo Fisher) blocking solution at room temperature for two hours. Each antibody was added to a primary antibody mix at the selected concentration in blocking solution and incubated overnight at 4°C. After two washes in PBS and PBS-0.1% Tween, the slides were treated with the DNA intercalator Cell-ID™ Intercalator-Ir (Fluidigm) (containing the two iridium isotopes 191Ir and 193Ir) 1.25 mM in a PBS solution. After a 30-minute incubation, the slides were washed once in PBS and once in MilliQ water and air-dried. The stained slides were then loaded in the Hyperion Imaging System (Fluidigm) imaging module to obtain light-contrast high resolution images of approximately four mm^2^. These images were used to select the ROI in each slide. For CLN1-CLN6, 1 mm^2^ ROIs were selected to contain the full thickness of the colon mucosa, with epithelial crypts in longitudinal orientation. For APP1, a one mm^2^ ROI containing a lymphoid follicle in its whole depth alongside a portion of lamina propria and of epithelium was selected. ROIs were ablated at a o µm/pixel resolution and 200 Hz frequency.

### IMC data analysis of human colon mucosa

Twenty-eight images from 26 antibodies (Supplementary Table 2) and two DNA intercalators were obtained from the raw .txt files of the ablated regions in CLN1-CLN6 using the data extraction process. Pixel intensities for each channel were normalised to the 99^th^ percentile in all samples and Otsu thresholding was performed on the normalized images with a custom CellProfiler4 pipeline, which was employed also to generate the masks for the lamina propria (using the Vimentin channel including all <75-pixel large negative areas) and the epithelium (starting from the Pan-keratin and E-cadherin channels, dilatating the images with a three-pixel disk and the filling up of all <75-pixel large negative areas). These masks were then added into a sum image, which underwent dilatation with a three-pixel disk and filling up of all <25-pixel large negative areas. Positive features outside of the lamina and epithelium were removed with an opening operation using a 150-pixel radius and the lamina propria mask was subtracted from the sum image to generate the final mask for the epithelial compartment. These masks and the thresholded images were used as input for the pixel-based and cell-based analysis processes. The IgA masks employed for the pixel analysis were generated using a three-class global Otsu thresholding with two background classes after applying a Gaussian filter with a 1.5-pixel large radius.

To evaluate the effect of normalisation on the downstream analysis, sample specific thresholds were manually selected for IgA, E-Cadherin, Pan-Keratin and Vimentin and applied to the raw images. The resulting thresholded images were used to generate lamina propria and epithelial masks for each sample individually.

Pixel-level analysis was performed on the IgA masks derived from either the normalised or the raw images and IgA^+^ areas in the tissue, lamina propria and epithelium were measured and normalised over the areas of the three compartments.

Cell-level analysis started with CellProfiler4 segmentation first on DNA1 with global Otsu thresholding to identify the cell nuclei. Then, cells were identified by radially expanding each nucleus for up to 10 pixels over a membrane mask derived from the IgA, CD3, CD68, CD11c and E-cadherin channels. After inspection by an expert histologist (JS), only cells overlapping with the lamina propria mask by at least 30% were retained.

Cell identities were assigned according to the highest overlap of the cell area with marker-specific thresholds defined by an expert histologist (JS): ≥ mask for IgA cells; ≥ of the CD3 mask for T cells; ≥ of the CD68 mask for macrophages; ≥15% of CD11c mask for dendritic cells.

### IMC data analysis of human appendix

Images from the same 26 antibodies and two DNA intercalators used in the colon mucosa (Supplementary Table 2) were obtained from the raw .txt files of the ablated region in APP1, normalised to the 99^th^ percentile and thresholded with CellProfiler4 as described above. For the cell-based analysis, nuclei were identified using the DNA1 channel and cells were isolated through watershed segmentation with the nuclei as seeds on a membrane mask summing up CD45, Pan-keratin and E-cadherin thresholded images.

Cells were assigned to the epithelium or to immune cell populations if they overlapped for ≥10% with the following masks: CD3 mask for T cells; CD20 and CD27 masks for B cells; CD68 mask for macrophages; CD11c mask for dendritic cells; E-cadherin^+^ and Pan-keratin^+^ masks for epithelial cells.

T cells were further phenotyped using unsupervised clustering at resolutions between 0.1 and 1.0, with 0.05 intervals and based on the cell marker intensity for CD3, CD45RA, CD45RO, CD4, CD8, Ki67 and PD1. The resulting clusters were manually inspected and the clustering with the highest number of biologically meaningful clusters (resolution = 0.25) was chosen. Clusters were re-identified using mean intensity thresholds defined by an expert histologist (JS) for the following markers: CD3 >0.06 for cluster 1; CD8a >0.125 for cluster 2; CD45RA >0.125 for cluster 3; CD4 >0.125 and CD45RO >0.15 for cluster 4; and CD4 > 0.1 and PD1 >0.15 for cluster 5.

Homotypic aggregations of CD4^+^PD1^+^ T cells (cluster 5, resolution = 0.25) were computed using a minimum of five points per cluster and a reachability parameter corresponding to a density of at least 5 cells/mm^2^.

### CD3 staining and mIF of human rectal cancer

Two 4 µm thick serial sections were cut from CRC1 FFPE block using a microtome. The first slide was dewaxed and rehydrated before carrying out HIER with Antigen Retrieval Reagent-Basic (R&D Systems). The tissue was then blocked and incubated with the anti-CD3 antibody (Dako, Supplementary Table 2) followed by horseradish peroxidase (HRP) conjugated anti-rabbit antibody (Dako) and stained with 3,3’ diaminobenzidine (DAB) substrate (Abcam) and haematoxylin. Areas with CD3^+^ infiltration in the proximity of the tumour invasive margin were identified by a clinical pathologist (M. R-J.)

The second slide was stained with a panel of six antibodies (CD8, PD1, Ki67, PDL1, CD68, GzB, Supplementary Table 2), Opal fluorophores and 4’,6-diamidino-2-phenylindole (DAPI) on a Ventana Discovery Ultra automated staining platform (Roche). Expected expression and cellular localisation of each marker as well as fluorophore brightness were used to minimise fluorescence spillage upon antibody-Opal pairing. Following a one-hour incubation at a 60°C, the slide was subjected to an automated staining protocol on an autostainer. The protocol involved deparaffinisation (EZ-Prep solution, Roche), HIER (DISC. CC1 solution, Roche) and seven sequential rounds of: one hour incubation with the primary antibody, 12 minutes incubation with the HRP-conjugated secondary antibody (DISC. Omnimap anti-Ms HRP RUO or DISC. Omnimap anti-Rb HRP RUO, Roche) and 16 minute incubation with the Opal reactive fluorophore (Akoya Biosciences). For the last round of staining, the slide was incubated with Opal TSA-DIG reagent (Akoya Biosciences) for 12 minutes followed by Opal 780 reactive fluorophore for our hour (Akoya Biosciences). A denaturation step (100°C for 8 minutes) was introduced between each staining round in order to remove the primary and secondary antibodies from the previous cycle without disrupting the fluorescent signal. The slide was counterstained with DAPI (Akoya Biosciences) and coverslipped using ProLong Gold antifade mounting media (Thermo Fisher Scientific). The Vectra Polaris automated quantitative pathology imaging system (Akoya Biosciences) was used to scan the labelled slide. Six fields of view, within the area selected by the pathologist, were scanned at 20x and 40x magnification using appropriate exposure times and loaded into inForm^41^ for spectral unmixing and autofluorescence isolation using the spectral libraries.

### mIF data analysis

After spectral unmixing and merging of six 20x fields of view for a total of >5mm^2^ ROI (Table 2), one single-tiff image was extracted for each marker and its intensity was rescaled from 0 to 1 with custom R scripts. The resulting single-tiff images were pre-processed to remove the background noise with Otsu thresholding in CellProfiler4 and used for cell segmentation by applying a global threshold to the DAPI channel and selecting all objects with a diameter between four and 60 pixels.

PD1^+^CD8^+^ cells, CD68^+^ cells and PDL1^+^ cells were then identified using mean intensity thresholds of 0.01 for CD8, 0.005 for PD1, 0.01 for CD68 and 0.01 for PDL1. All thresholds were inspected by an expert histologist (JS).

The distributions of minimum distances between PDL1^+^ cells and PD1^+^CD8^+^ cells were calculated from the coordinates of the centroids of each cell in the image. All PDL1^+^ cells and PD1^+^CD8^+^ cells at a distance from each other lower than double the maximum cell radius (24 pixels = 12 µm) were considered as proximal. All other cells were classified as distal.

### CODEX data analysis

A published dataset of colorectal CODEX images^9^ was downloaded from The Cancer Imaging Archive (https://doi.org/10.7937/tcia.2020.fqn0-0326). It consisted of processed CODEX data from 35 colorectal cancer samples divided in two groups (CLR and DII) according to the peritumoral inflammatory levels and the presence of tertiary lymphoid structures^9^. For each sample, four .tiff images were available representing four 0.6mm spots from two 70-core tissue microarrays. These images were hyperstacks of 58 channels including 56 antibodies (Supplementary Table 2) and two DNA markers with a resolution of 377 nm/pixel. After manual review of all 140 spots, one representative image per sample was selected, having the best focus and containing both tumour and peritumoural immune infiltrates.

The single-channel tiff files for each selected image were extracted and the pixel intensities were rescaled from 0 to 1 with a custom R script. Using SIMPLI, single-cell segmentation was performed in each of the 35 images by applying a global threshold to the HOECHST channel to identify the nuclei and retaining all objects with a diameter between 5 and 40 pixels. Each nucleus was then expanded by 5 pixels in all directions to define the cell area.

Resulting single cells were assigned to ten phenotypes according to the mean cell expression of CDX2 >0.15 or MUC1 >0.15 or cytokeratin >0.15 for tumour cells; CD34 >0.15 or CD31 >0.15 for endothelial cells; vimentin >0.1 for other stromal cells; CD11c >0.3 for dendritic cells; CD38 >0.26 for B cells; CD4 >0.13 and CD3 >0.1 for CD4^+^ T cells; CD4 >0.12 and FOXP3 >0.5 and CD3 >0.1 for Tregs; CD8 >0.16 and CD3 >0.1 for CD8^+^ T cells, and CD68 >0.11 for macrophages. The heterotypic spatial analysis was performed by calculating the minimum distances of macrophages, CD8^+^ T cells, CD4^+^ T cells, Treg cells, and B cells to tumour cells and endothelial cells using the coordinates of the cell centroids.

## DATA AVAILABILITY

The data that support the findings of this study are available from the corresponding author, F.D.C., upon request.

## CODE AVAILABILITY

SIMPLI’s code, documentation and an example dataset are available at https://github.com/ciccalab/SIMPLI. The software code is protected by copyright. No permission is required from the rights-holder for non-commercial research uses. Commercial use will require a license from the rights-holder. For further information contact.

## ACKNOWLEDGMENTS

We thank Sharavan Vishaan Venkateswaran for testing SIMPLI. F.C is supported by Cancer Research UK [C43634/A25487], the Cancer Research UK King’s Health Partners Centre at King’s College London [C604/A25135], the Cancer Research UK City of London Centre [C7893/A26233], innovation programme under the Marie Skłodowska-Curie grant agreement No CONTRA-766030 and the Francis Crick Institute, which receives its core funding from Cancer Research UK (FC001002), the UK Medical Research Council (FC001002), and the Wellcome Trust (FC001002). For the purpose of Open Access, the author has applied a CC BY public copyright licence to any Author Accepted Manuscript version arising from this submission.

## AUTHOR CONTRIBUTIONS

F.D.C. conceived and directed the study with support of J.S. M.B. developed the software with the help of D.T. and M.R.K. L.M. and A.A.S performed the experiments. M.B, L.M, A.A.S., M.J.P., J.S., and F.D.C. analysed the data. M.R.J, G.B. and L.L. identified the samples and provided clinical assessments. M.R.J. performed pathological assessments. M.B. and F.D.C. wrote the manuscript with contribution from A.A.S., L.M. and M.J.P. All authors approved the manuscript.

## REFERENCES

1. Parra ER, Francisco-Cruz A, Wistuba II. State-of-the-art of profiling immune contexture in the era of multiplexed staining and digital analysis to study paraffin tumor tissues. Cancers 11, 247 (2019).

2. Giesen C, et al. Highly multiplexed imaging of tumor tissues with subcellular resolution by mass cytometry. Nature Methods 11, 417–422 (2014).

3. Angelo M, et al. Multiplexed ion beam imaging of human breast tumors. Nature medicine 20, 436–442 (2014).

4. Goltsev Y, et al. Deep Profiling of Mouse Splenic Architecture with CODEX Multiplexed Imaging. Cell 174, 968–981.e915 (2018).

5. Lin JR, Fallahi-Sichani M, Chen JY, Sorger PK. Cyclic immunofluorescence (CycIF), a highly multiplexed method for single-cell imaging. Current protocols in chemical biology 8, 251–264 (2016).

6. Au - Bauman TM, Au - Ricke EA, Au - Drew SA, Au - Huang W, Au - Ricke WA. Quantitation of Protein Expression and Co-localization Using Multiplexed Immuno-histochemical Staining and Multispectral Imaging. JoVE, e53837 (2016).

7. Morrison LE, et al. Brightfield multiplex immunohistochemistry with multispectral imaging. Laboratory Investigation 100, 1124–1136 (2020).

8. Jackson HW, et al. The single-cell pathology landscape of breast cancer. Nature 578, 615–620 (2020).

9. Schürch CM, et al. Coordinated Cellular Neighborhoods Orchestrate Antitumoral Immunity at the Colorectal Cancer Invasive Front. Cell 182, 1341–1359.e1319 (2020).

10. Berg S, et al. ilastik: interactive machine learning for (bio)image analysis. Nature Methods 16, 1226–1232 (2019).

11. McQuin C, et al. CellProfiler 3.0: Next-generation image processing for biology. PLOS Biology 16, e2005970 (2018).

12. van Maldegem F, et al. Characterisation of tumour immune microenvironment remodelling following oncogene inhibition in preclinical studies using an optimised imaging mass cytometry workflow. bioRxiv, 2021.2002.2002.429358 (2021).

13. Zanotelli VRT, Bodenmiller B. ImcSegmentationPipeline: A pixelclassification based multiplexed image segmentation pipeline. Zenodo, (2017).

14. Jones TR, et al. CellProfiler Analyst: data exploration and analysis software for complex image-based screens. BMC Bioinformatics 9, 482 (2008).

15. Eling N, Damond N, Hoch T, Bodenmiller B. cytomapper: an R/Bioconductor package for visualisation of highly multiplexed imaging data. bioRxiv, 2020.2009.2008.287516 (2020).

16. Opzoomer JW, et al. ImmunoCluster provides a computational framework for the nonspecialist to profile high-dimensional cytometry data. eLife 10, e62915 (2021).

17. Stoltzfus CR, et al. CytoMAP: A Spatial Analysis Toolbox Reveals Features of Myeloid Cell Organization in Lymphoid Tissues. Cell Reports 31, 107523 (2020).

18. Somarakis A, Unen VV, Koning F, Lelieveldt B, Höllt T. ImaCytE: Visual Exploration of Cellular Micro-Environments for Imaging Mass Cytometry Data. IEEE Transactions on Visualization and Computer Graphics 27, 98–110 (2021).

19. Yang T, et al. SPIAT: An R package for the Spatial Image Analysis of Cells in Tissues. bioRxiv, 2020.2005.2028.122614 (2020).

20. neighbouRhood. https://githubcom/BodenmillerGroup/neighbouRhood, (2019).

21. Catena R, Montuenga LM, Bodenmiller B. Ruthenium counterstaining for imaging mass cytometry. The Journal of pathology 244, 479–484 (2018).

22. Bankhead P, et al. QuPath: Open source software for digital pathology image analysis. Scientific Reports 7, 16878 (2017).

23. Dries R, et al. Giotto: a toolbox for integrative analysis and visualization of spatial expression data. Genome Biology 22, 78 (2021).

24. Schmidt U, Weigert M, Broaddus C, Myers G. Cell Detection with Star-Convex Polygons. (ed^(eds). Springer International Publishing (2018).

25. Pabst O, Slack E. IgA and the intestinal microbiota: the importance of being specific. Mucosal Immunology 13, 12–21 (2020).

26. Dorn I, Schlenke P, Mascher B, Stange EF, Seyfarth M. Lamina Propria Plasma Cells in Inflammatory Bowel Disease: Intracellular Detection of Immunoglobulins Using Flow Cytometry. Immunobiology 206, 546–557 (2002).

27. Song W, Craft J. T follicular helper cell heterogeneity: Time, space, and function. Immunological Reviews 288, 85–96 (2019).

28. Zhang L, et al. Lineage tracking reveals dynamic relationships of T cells in colorectal cancer. Nature 564, 268–272 (2018).

29. Strokotov D, et al. Is there a difference between T- and B-lymphocyte morphology? Journal of Biomedical Optics 14, 064036 (2009).

30. imctools. https://githubcom/BodenmillerGroup/imctools, (2017).

31. Pau G, Fuchs F, Sklyar O, Boutros M, Huber W. EBImage—an R package for image processing with applications to cellular phenotypes. Bioinformatics 26, 979–981 (2010).

32. Van der Walt S, et al. scikit-image: image processing in Python. PeerJ 2, e453 (2014).

33. Butler A, Hoffman P, Smibert P, Papalexi E, Satija R. Integrating single-cell transcriptomic data across different conditions, technologies, and species. Nature Biotechnology 36, 411–420 (2018).

34. Blondel VD, Guillaume J-L, Lambiotte R, Lefebvre E. Fast unfolding of communities in large networks. Journal of statistical mechanics: theory and experiment 2008, P10008 (2008).

35. Leland M, John H, Nathaniel S, Lukas G. UMAP: uniform manifold approximation and projection. Journal of Open Source Software 3, 861 (2018).

36. Ester M, Kriegel H-P, Sander J, Xu X. A density-based algorithm for discovering clusters in large spatial databases with noise. In: Kdd (ed^(eds) (1996).

37. Henning C. fpc. https://cranr-projectorg/web/packages/fpc/indexhtml, (2020).

38. Di Tommaso P, Chatzou M, Floden EW, Barja PP, Palumbo E, Notredame C. Nextflow enables reproducible computational workflows. Nature Biotechnology 35, 316–319 (2017).

39. Kurtzer GM, Sochat V, Bauer MW. Singularity: Scientific containers for mobility of compute. PLOS ONE 12, e0177459 (2017).

40. Sochat VV, Prybol CJ, Kurtzer GM. Enhancing reproducibility in scientific computing: Metrics and registry for Singularity containers. PLOS ONE 12, e0188511 (2017).

41. Kramer AS, et al. InForm software: a semi-automated research tool to identify presumptive human hepatic progenitor cells, and other histological features of pathological significance. Scientific Reports 8, 3418 (2018).

