## Supplementary figures for "A SIMPLI (Single-cell Identification from MultiPLexed Images) approach for spatially resolved tissue phenotyping at single-cell resolution"

**Supplementary Figure 1.** SIMPLI workflow diagram

**Supplementary Figure 2.** Pixel analysis and cell masking of human colon mucosa

**Supplementary Figure 3.** Comparison of T cell phenotypes in human appendix

**Supplementary Table 1.** Samples used in the study

**Supplementary Table 2.** Antibodies used in the study

**Supplementary Figure 1. SIMPLI workflow diagram**

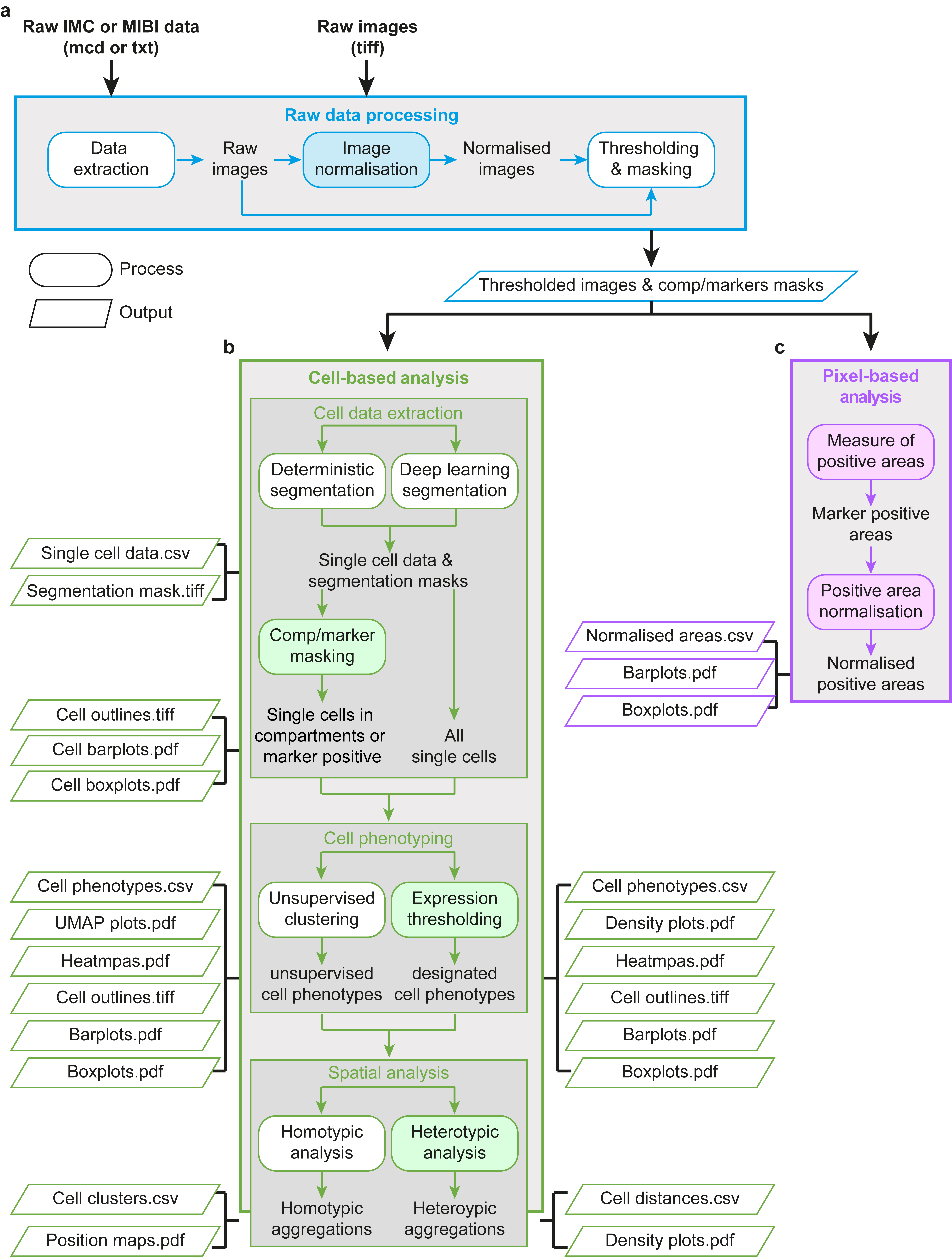

SIMPLI’s workflow is divided into three main steps: raw data processing (**a**), cell-based analysis (**b**) and pixel-based analysis (**c**). Each step is divided in multiple stand-alone processes (rectangles), which rely on established tools and libraries (white) or newly developed codes (blue, green and pink), and produces multiple outputs (parallelograms).

**a.** Raw data processing. Raw data from IMC or MIBI (.mcd or .txt) are extracted using imctools^1^. Resulting images or original .tiff images from other imaging platforms are normalised using custom scripts and thresholded with a containerised headless instance of CellProfiler^2^ to produce tissue compartments or marker masks as well as images for the following steps.

**b.** Cell-based analysis. This step is divided into cell data extraction, cell phenotyping and spatial analysis. Single cells are identified through single-cell segmentation using CellProfiler^2^ or StarDist^3^ with default or user-provided trained models. Cells belonging to tissue compartments or positive for certain markers can be identified based on their overlap with the tissue compartments or marker masks derived in the previous step. Subsequently, cell phenotypes are refined using unsupervised clustering with Seurat^4^ or applying expression thresholds to one or more markers using *ad hoc* scripts. Finally, the spatial distribution of homotypic cell aggregations is performed with DBSCAN^5^, while heterotypic cell aggregations are investigated using custom scripts.

**c.** Pixel-based analysis. Areas positive for a specific marker or combination of markers are measured from the thresholded images and normalised over the area of the whole image or tissue compartments. These normalised values can then be compared across datasets. All processes in this step are performed using *ad hoc* scripts integrated in the pipeline.

**Supplementary Figure 2.** Pixel analysis and cell masking of human colon mucosa

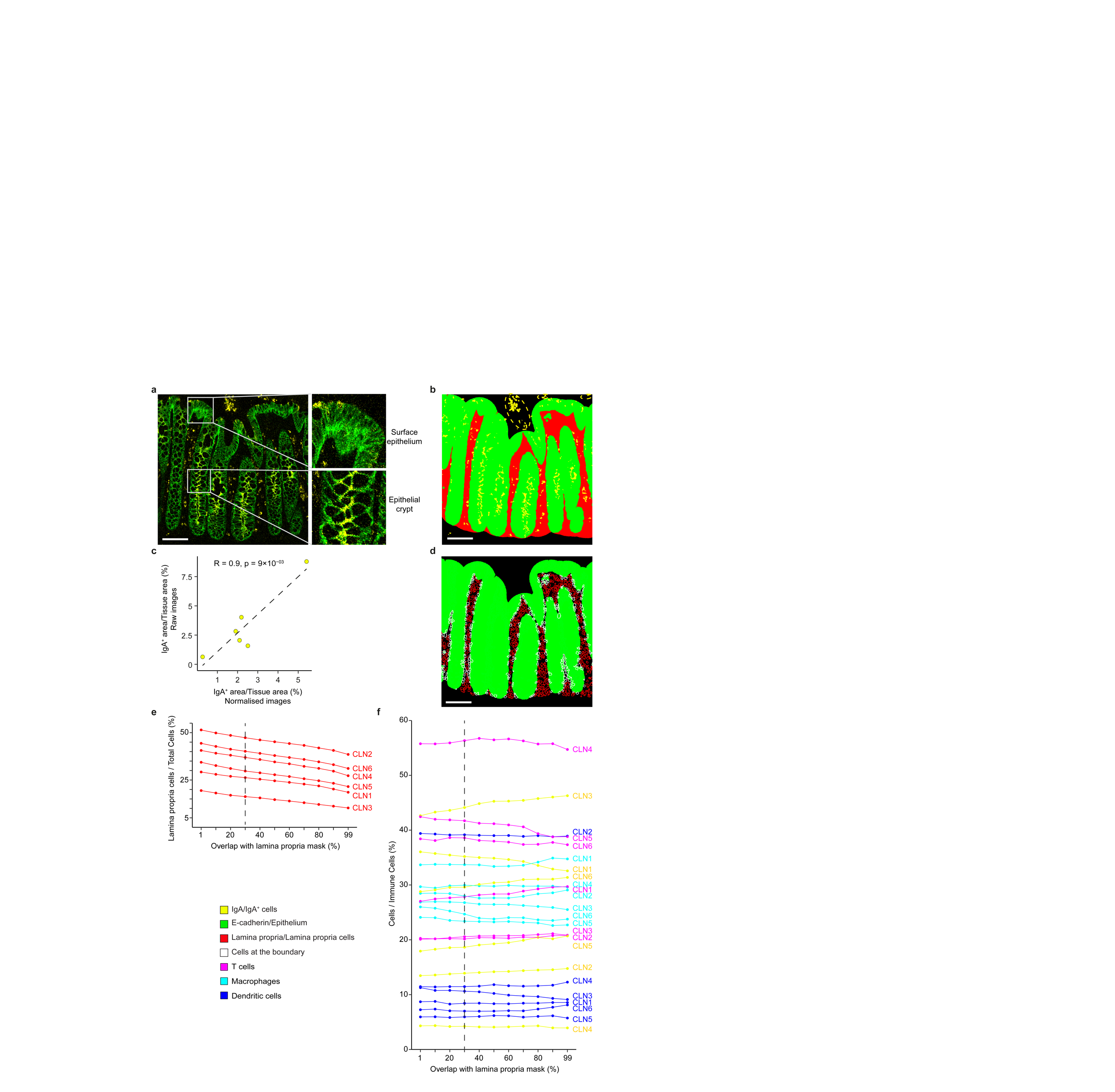

**a.** IMC image of normal colon mucosa in CLN6 after data extraction and normalisation. Zoom-ins illustrate examples of surface epithelium and epithelial crypts. IgA^+^ pixels are concentrated in the epithelial crypts where most of IgA transcytosis takes place.

**b.** Distribution of IgA^+^ pixels in CLN6. Epithelium and lamina propria masks were generated as described in the Methods and superimposed to the mask of the IgA channel. Only IgA^+^ pixels within the two compartments were retained for the pixel analysis, thus excluding likely artefacts (dotted circles). Scale bar in (a) and (b) = 100μm.

**c.** Correlation between IgA^+^ areas measured from raw and normalised images. Pearson correlation coefficient R and associated p-value are shown.

**d.** Cells at the boundary between epithelium and lamina propria in CLN6. These were defined as cells with a partial overlap with both masks and their assignment to either compartment depends on the overlap threshold.

Parallel plots of all cells (**e**) and only immune cells (**f**) resident in the lamina propria at various thresholds of overlap (1% to 99% of the total cell area). Dotted lines represent the value chosen for the downstream analysis in Figure 2e,f (30%).

**Supplementary Figure 3.** Comparison of T cell phenotypes in human appendix

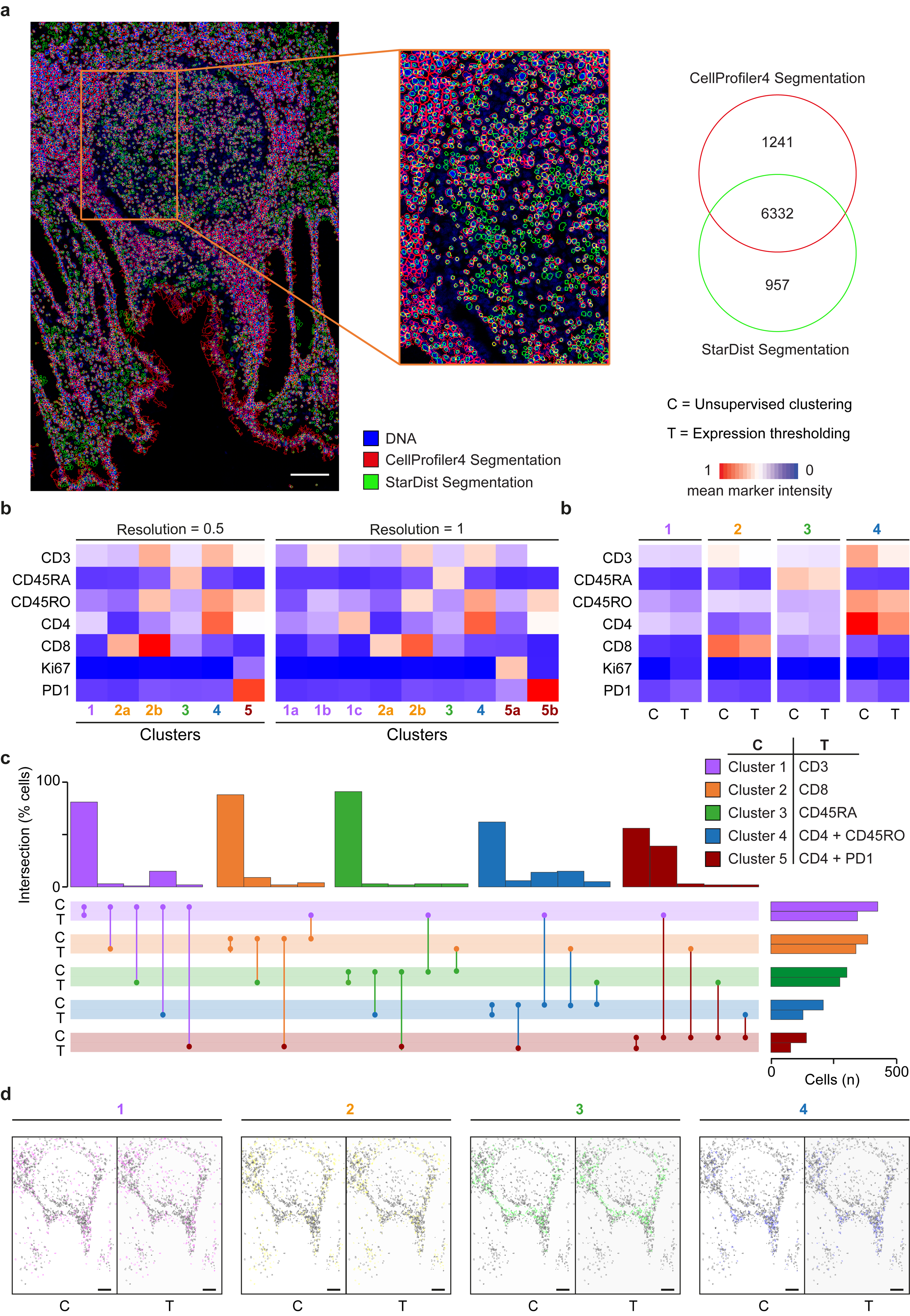

**a.** Comparison of single-cell segmentations of APP1 obtained with CellProfiler4 and StarDist superimposed over the normalised DNA masks. The two segmentations were performed as described in the Methods leading to the majority of cells identified by both approaches.

**b.** Expression profiles of T cell subpopulations identified using unsupervised clustering with resolution of 0.5 and 1.0. Clusters are numbered as in Figure 3e showing how increasing the resolution splits bigger clusters obtained at lower resolution.

**c.** Comparison of the expression profiles of T cell subpopulations identified using unsupervised clustering at 0.25 resolution and expression thresholding of representative markers. For each population in (**b**) and (**c**), the mean intensity value of the markers across all cells is shown. The colour scale was normalised across all markers and cells, independently for each analysis.

**d.** Percentage of cells shared between clustering-derived and thresholding-derived phenotypes. Number of cells identified by the two classification methods in each population are also reported in the lateral bars.

**e.** Position map of T cells in APP1 colour-coded according to the phenotype obtained through unsupervised clustering or expression thresholding. Scale bar = 100μm.

**References**

1.	imctools. [*https://githubcom/BodenmillerGroup/imctools*](https://githubcom/BodenmillerGroup/imctools), (2017).

2.	McQuin C*, et al.* CellProfiler 3.0: Next-generation image processing for biology. *PLOS Biology* **16**, e2005970 (2018).

3.	Schmidt U, Weigert M, Broaddus C, Myers G. Cell Detection with Star-Convex Polygons. (ed^(eds). Springer International Publishing (2018).

4.	Butler A, Hoffman P, Smibert P, Papalexi E, Satija R. Integrating single-cell transcriptomic data across different conditions, technologies, and species. *Nature Biotechnology* **36**, 411-420 (2018).

5.	Ester M, Kriegel H-P, Sander J, Xu X. A density-based algorithm for discovering clusters in large spatial databases with noise. In: *Kdd* (ed^(eds) (1996).
